# Phylogenomics-guided discovery of a novel conserved cassette of short linear motifs in BubR1 essential for the spindle checkpoint

**DOI:** 10.1101/088161

**Authors:** Eelco Tromer, Debora Bade, Berend Snel, Geert J.P.L. Kops

## Abstract

The spindle assembly checkpoint (SAC) maintains genomic integrity by preventing progression of mitotic cell division until all chromosomes are stably attached to spindle microtubules. The SAC critically relies on the paralogs Bub1 and BubR1/Mad3, which integrate kinetochore-spindle attachment status with generation of the anaphase inhibitory complex MCC. We previously reported on the widespread occurrences of independent gene duplications of an ancestral ‘MadBub’ gene in eukaryotic evolution and the striking parallel subfunctionalization that lead to loss of kinase function in BubR1/Mad3-like paralogs. Here, we present an elaborate subfunctionalization analysis of Bub1/BubR1 gene family and perform de novo sequence discovery in a comparative phylogenomics framework to trace the distribution of ancestral sequence features to extant paralogs throughout the eukaryotic tree of life. We show that known ancestral sequence features are consistently retained in the same functional paralog: GLEBS/CDI/CDII/kinase in the Bub1-like and KEN1/KEN2/D-Box in the BubR1/Mad3-like. The recently described ABBA motif can be found in either or both paralogs. We however discovered two additional ABBA motifs that flank KEN2. This cassette of ABBA1-KEN2-ABBA2 forms a strictly conserved module in all ancestral and BubR1/Mad3-like proteins, suggestive of a specific and crucial SAC function. Indeed, deletion of the ABBA motifs in human BUBR1 abrogates the SAC and affects APC/CCdc20 interactions. Our detailed comparative genomics analyses thus enabled discovery of a conserved cassette of motifs essential for the SAC and shows how this approach can be used to uncover hitherto unrecognized functional protein features.

## Introduction

Chromosome segregation during cell divisions in animals and fungi is monitored by a cell cycle checkpoint known as the spindle assembly checkpoint (SAC) (Vleugel et al., 2012; London and Biggins, 2014a; Musacchio, 2015). The SAC couples absence of stable attachments between kinetochores and spindle microtubules to inhibition of anaphase by assembling a four-subunit inhibitor of the anaphase-promoting complex (APC/C), known as the MCC (Sacristan and Kops, 2015; Izawa and Pines, 2014; Chao et al., 2012). The molecular pathway that senses lack of attachment and produces the MCC relies on two related proteins known as Bub1 and BubR1/Mad3 (London and Biggins, 2014a). Bub1 is a serine/threonine kinase that localizes to kinetochores and promotes recruitment of MCC subunits and of factors that stimulate its assembly (Klebig et al., 2009; London and Biggins, 2014b; Vleugel et al., 2015) These events are largely independent of Bub1 kinase activity, however, which instead is essential for the correction process of attachment errors (Kawashima et al., 2010; Klebig et al., 2009; Andrews et al., 2004). BubR1/Mad3 is one of the MCC subunits, responsible for directly preventing APC/C activity and anaphase onset (Tang et al., 2001; Sudakin et al., 2001; Chao et al., 2012). It does so by contacting multiple molecules of the APC/C co-activator Cdc20, preventing APC/C substrate access and binding of the E2 enzyme UbcH10 (Chao et al., 2012; Izawa and Pines, 2014; Alfieri et al., 2016; Yamaguchi et al., 2016). The BubR1/Mad3-Cdc20 contacts occur via various short linear motifs (SLiMs) known as ABBA, KEN, and D-box (Alfieri et al., 2016; Chao et al., 2012; Lischetti et al., 2014; King et al., 2007; DiFiore et al., 2015; Burton and Solomon, 2007b). Like Bub1, BubR1 also impacts on the attachment error-correction process via a KARD motif that recruits the PP2A-B56 phosphatase (Suijkerbuijk et al., 2012b; Kruse et al., 2013; Xu et al., 2013). This may not however be a universal feature of BubR1/Mad3-like proteins, because many lack a KARD-like motif.

Bub1 and BubR1/Mad3 are paralogs. We previously showed they originated by similar but independent gene duplications from an ancestral MadBub gene in many lineages, and that the two resulting gene copies then subfunctionalized in remarkably comparable ways (Suijkerbuijk et al., 2012a). An ancestral N-terminal KEN motif (KEN1: essential for the SAC) and an ancestral C-terminal kinase domain (essential for attachment error-correction) were retained in only one of the paralogous genes in a mutually exclusive manner in virtually all lineages (i.e. one gene retained KEN but lost kinase, while the other retained kinase but lost KEN). One exception to this ‘rule’ are vertebrates where both paralogs have a kinase-like domain. The kinase domain of human BUBR1 however lacks enzymatic activity (i.e. is a pseudokinase) but instead confers stability onto the BUBR1 protein (Suijkerbuijk et al., 2012a).

The similar subfunctionalization of Bub1 and BubR1/Mad3-like paralogs was inferred from analysis of two domains (TPR and kinase) and one motif (KEN1). We set out to analyze whether any additional features specifically segregated to Bub1‐ or BubR1/Mad3-like proteins after duplications by designing an unbiased feature discovery pipeline and tracing feature evolution. The pipeline extracted all known and various previously unrecognized conserved motifs from Bub1/BubR1 family gene members. Two of these are novel ABBA motifs that flank KEN2 specifically in BuBr1/Mad3-like proteins and we show that this highly conserved and ABBA-KEN2-ABBA cassette is crucial for the SAC in human cells.

## Results and Discussion

### Refined phylogenomic analysis of the MadBub gene family pinpoints 16 independent gene duplication events across the eukaryotic tree of life

To enable detailed reconstruction of subfunctionalization events of all known functional features after duplication of ancestral MadBub genes, we expanded our previously published set of homologs (Suijkerbuijk et al., 2012a) through broader sampling of sequenced eukaryotic genomes, focusing on sequences closely associated with duplication events (**supplementary sequence file 1**). Phylogenetic analyses of a multiple sequence alignment of the TPR domain (the only domain shared by all MadBub family members) of 149 MadBub homologs (supplementary discussion and supplementary figure 1) corroborated the ten independent duplications previously described (Suijkerbuijk et al., 2012a) and allowed for a more precise determination of the age of the duplications. Strikingly, we found evidence for a number of additional independent duplications: Three duplications in stramenopile species of the SAR super group (Albuginaceae {#10 in **figure 1b**}, E. siliculosis {#11} and A. anophagefferens {#12}) and one at the base of basidiomycete fungi ({#4}, puccinioimycetes). The BUBR1 paralog in teleost fish underwent a duplication and fission event, of which the C-terminus product was retained only in the lineage leading to zebra fish (D. rerio {#7}). Lastly, through addition of recently sequenced genomes we could specify a duplication around the time plants started to colonize land ({#13}, bryophytes) and an independent duplication in the ancestor of higher plants ({#14}, tracheophytes), followed by a duplication in the ancestor of the flowering plants ({#15}, magnoliaphytes). These gave rise to three MadBub homologs, signifying additional subfunctionalization of the paralogs in the plant model organism A. thaliana. It thus seems to be the case that such striking parallel subfunctionalization as we originally identified, is indeed predictive for more of its occurrence in lineages whose genome sequences have since been elucidated.

**Figure 1.**
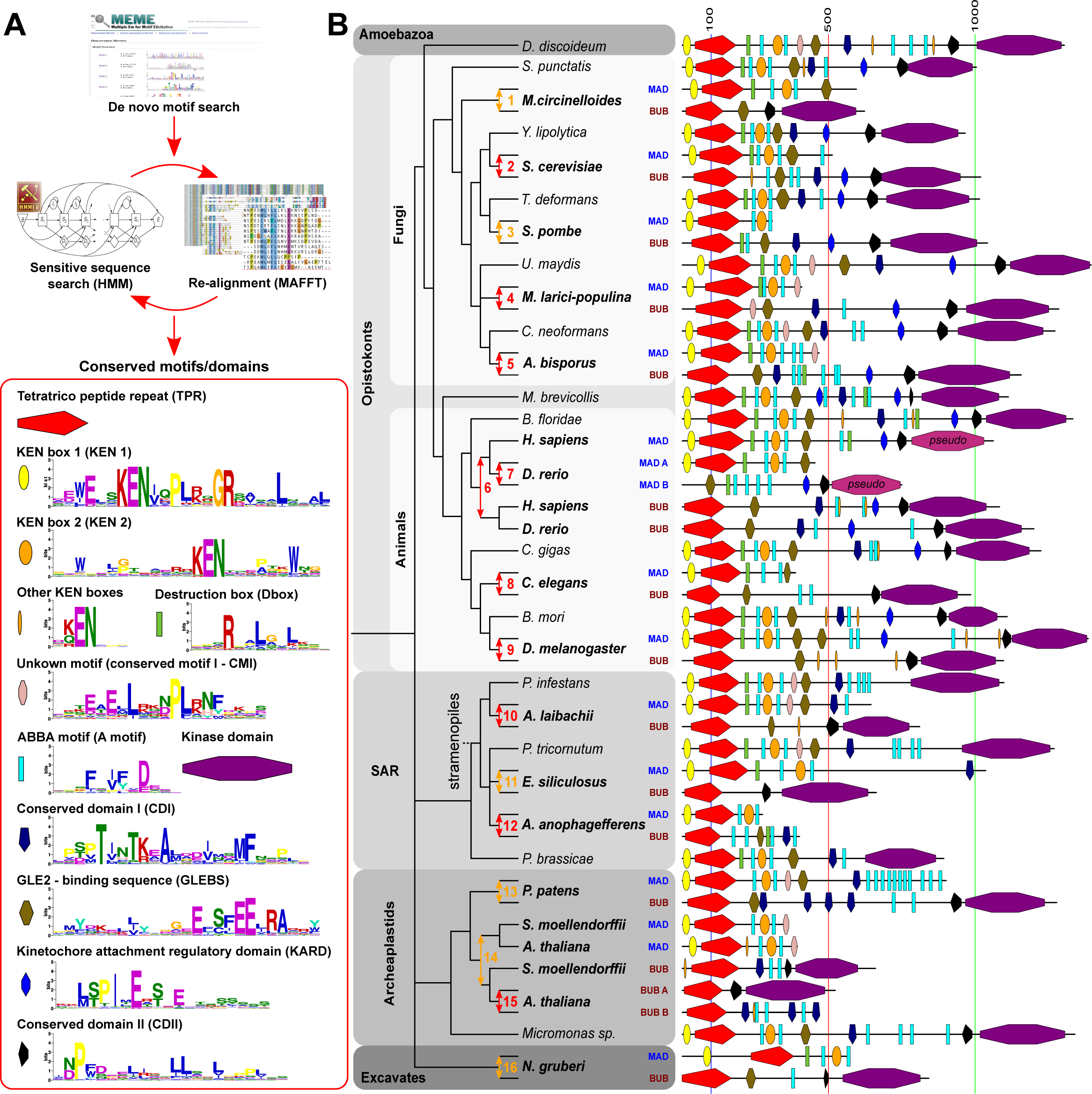
Fate of conserved functional sequence features after 16 independent duplications of the MADBUB gene family throughout eukaryotic evolution. (A) Overview of the de novo sequence discovery pipeline ConFeaX including the ancestral conserved features of a search against the eukaryotic MADBUB gene family. The consensus sequences of the detected conserved motifs are depicted as a sequence logo (colors reflect distinct amino acid properties and height of the letters indicates conservation of amino acids). Each features is assigned a differently colored shape. (B) Cartoon of the evolutionary scenario of 16 independent duplications of the MADBUB gene family throughout eukaryotic evolution, including a projection of conserved features onto the linear protein representation (on scale). Gene duplications are indicated by an arrow (red: high confidence, orange: uncertain). The subfunctionalized paralogs MAD and BUB are color brown and blue, respectively. Numbers indicate the clades in which the duplications occurred: {1}-mucorales; {2}-saccharomycetaceae; {3}-schizosaccharomycetes; {4}-pucciniomycetes: {5}-agaricomycetes (excluding earlybranching species); {6}-vertebrates; {7}-teleost fish; {8}-nematodes; {9}-diptera (flies); {10}-albuginaceae (oomycete); {11}-ectocarpales (brow algae); {12}-aureococcus (harmful algae bloom); {13}-bryophytes (mosses); {14}-tracheophytes (vascular plants); {15}-magnoliaphytes (flowering plants); {16}-naegleria;

### De novo discovery, phylogenetic distribution and fate after duplication of functional motifs in the MadBub gene family

Previous analyses revealed a recurrent pattern of mutually exclusive retention of an N-terminal KEN-box and a C-terminal kinase domain after duplication of an ancestral MadBub (Suijkerbuijk et al., 2012a; Murray, 2012). These patterns suggested the hypothesis of paralog subfunctionalization towards either inhibition of the APC/C in the cytosol (retaining the KEN-box) or attachment-error correction at the kinetochore (retaining the kinase domain). Given the extensive sequence divergence of MadBub homologs and a scala of different known functional elements, we reasoned that a comprehensive analysis of MadBub gene duplicates would provide opportunities for the discovery of novel and co-evolving ancestral features. For clarity we refer to the Bub1-like paralog (C-terminal kinase domain) as BUB and the BubR1/Mad3-like paralog (N-terminal KEN box) as MAD throughout the rest of this manuscript.

To capture conserved ancestral features of diverse eukaryotic MadBub homologs, we constructed a sensitive de novo motif and domain discovery pipeline (ConFeaX: conserved feature extraction) similar to our previous approach used to characterize KNL1 evolution (Tromer et al., 2015). In short, the MEME algorithm (Bailey et al., 2009) was used to search for significantly similar gapless amino acid motifs, and extended motifs were aligned by MAFFT (Katoh and Standley, 2013). Alignments were modeled using HMMER (Eddy, 2011) and sensitive profile HMM searches were iterated and specifically optimized using permissive E-values/bit scores until convergence (**methods** and **figure 1a**). Due to the degenerate nature of the detected short linear motifs, we manually scrutinized the results for incorrectly identified features and supplemented known motif instances, when applicable. We preferred ConFeaX over other de novo motif discovery methods (e.g. Davey et al., 2012 Nguyen Ba et al., 2012) as it does not rely on high quality full length alignment of protein sequences and allows detection of repeated or dynamic non-syntenic conserved features (which is a common feature for SLiMs). It is therefore better tuned to finding conserved features over long evolutionary distances in general and specifically in this case where recurrent duplication and subfunctionalization hamper conventional multiple sequence alignment based analysis.

ConFeaX identified known functional motifs and domains and in some cases extended their definition: KEN1 (Murray and Marks, 2001), KEN2 (Burton and Solomon, 2007a), GLEBS (Taylor et al., 1998), KARD (Suijkerbuijk et al., 2012b; Xu et al., 2013; Kruse et al., 2013), CDI (Klebig et al., 2009), D-box (Burton and Solomon, 2007a), CDII (a co-activator domain of BUB1 (Kang et al., 2008; Klebig et al., 2009)) and the recently discovered ABBA motif (termed ABBA3 in **figure 3**) (DiFiore et al., 2015; Lischetti et al., 2014) (**figure 1a**, **supplementary table II** and **supplementary sequence file 2**). The TPR and the kinase domain were annotated using profile searches of previously established models (Suijkerbuijk et al., 2012a) and excluded from de novo sequence searches. KEN1 and KEN2 could be discriminated by differentially conserved residues surrounding the core KEN box (**figure 1a**). Those surrounding KEN1 are involved in the formation of the helix-turn-helix motif that positions yeast Mad3 towards Cdc20 (Chao et al., 2012), while two pseudo-symmetrically conserved tryptophan residues with unknown function specifically defined KEN2. Furthermore, we found that the third position of the canonical ABBA motif is often occupied by a proline residue and the first position in ascomycetes (fungi) is often substituted for a polar amino acid [KRN] (**figure 1a**), signifying potential lineage-specific changes in Cdc20-ABBA interactions. Last, we also discovered a novel motif predominantly associated with the MAD paralog in basidiomycetes, plants, amoeba and stramenopiles but not metazoa, which we termed conserved motif I (CMI) (**figure 1a**).

Projection of the conserved ancestral features onto the MadBub gene phylogeny provided a highly detailed overview of MadBub motif evolution (**figure 1b**, **supplementary figure 1b**). We found that the core functional motifs and domains (TPR, KEN1, KEN2, ABBA, D-box, GLEBS, CMI, CDI, CDII and kinase) are present throughout the eukaryotic tree of life, representing the core features that were likely part of the SAC signaling network in the last eukaryotic common ancestor (LECA). Of note are lineages (nematodes, flatworms (S. mansoni), dinoflagellates (S. minutum) and early-branching fungi (microsporidia and C. coronatus)) for which multiple features were either lost or considerably divergent (**supplementary figure 1b**). Especially interesting is C. elegans in which both KEN boxes and the GLEBS domain appeared to have been degenerated (ceMAD=san-1) and the CDI domain is lost (ceBUB=bub-1), indicating extensive rewiring or a less essential role of the SAC in nematode species, as has been suggested recently (Davey and Morgan, 2016; Moyle et al., 2014).

Our motif discovery analyses revealed the CDC20/Cdh1-interacting ABBA motif to be much more abundant than the single instances that were previously reported for BUBR1 and BUB1 in humans (Lischetti et al., 2014; DiFiore et al., 2015). We observed three different contexts for the ABBA motifs (**figure 1b**, **supplementary figure 1b**): (1) in repeat arrays (e.g. MAD of P. patens, basidiomycetes and stramenopiles), (2) in the vicinity of CDI (many instances) and/or D-box/KEN (e.g. human), and (3) as two highly conserved ABBA motifs flanking KEN2 (virtually all species). Because of the positional conservation of the latter, we have termed these, ABBA1 and ABBA2. Any additional ABBA motifs were pooled in the category ‘ABBA-other’.

In order to track the fate of the features discovered using ConFeaX, we quantified their co-presences and absences, as a proxy for co-evolution, by calculating the Pearson correlation coefficient (r) for the profiles of each domain/motif pair of 16 duplicated MadBub homologs (**figure 1b**)(Wu et al., 2003). Subsequent average clustering of the Pearson distance (d=1-r) revealed two sets of co-segregating and anti-correlated conserved features (**figure 2a+b**) consistent with our hypothesis that MadBub gene duplication caused parallel subfunctionalization of features towards the kinetochore (mainly BUB) and the cytosol (MAD) (Suijkerbuijk et al., 2012a). GLEBS, CDI, ABBA-other, KARD, CDII and the kinase domain formed a coherent cluster of features with bona fide function at the kinetochore. For a detailed discussion on several intriguing observations regarding presence/absence of these motifs in several eukaryotic lineages, and what this may mean for BUB/MAD and SAC function in these lineages, see **Supplementary Discussion**. A second cluster contained known motifs that bind and interact with (multiple) CDC20 molecules, including KEN1, KEN2, and (to a lesser extent) the D-box. Our newly discovered ABBA motifs that flank KEN2 were tightly associated with KEN2 and KEN1 (**figure 2**). As such, the ABBA1-KEN2-ABBA2 cassette (**Figure 3a**) co-segregated with MAD function during subfunctionalization of MadBub gene duplicates. Although the D-box often co-occurs with the KEN-ABBA cluster, this motif was occasionally lost (e.g. archeaplastids, S. pombe and A. anophagefferens). Finally, CMI co-segregated with the Cdc20-interacting motifs (**Figure 2a**), suggesting a MAD-specific role for this newly discovered motif (possibly in MCC function and/or Cdc20-binding) in species harboring it such as plants, basidiomycetes and stramenopiles.

**Figure 2.**
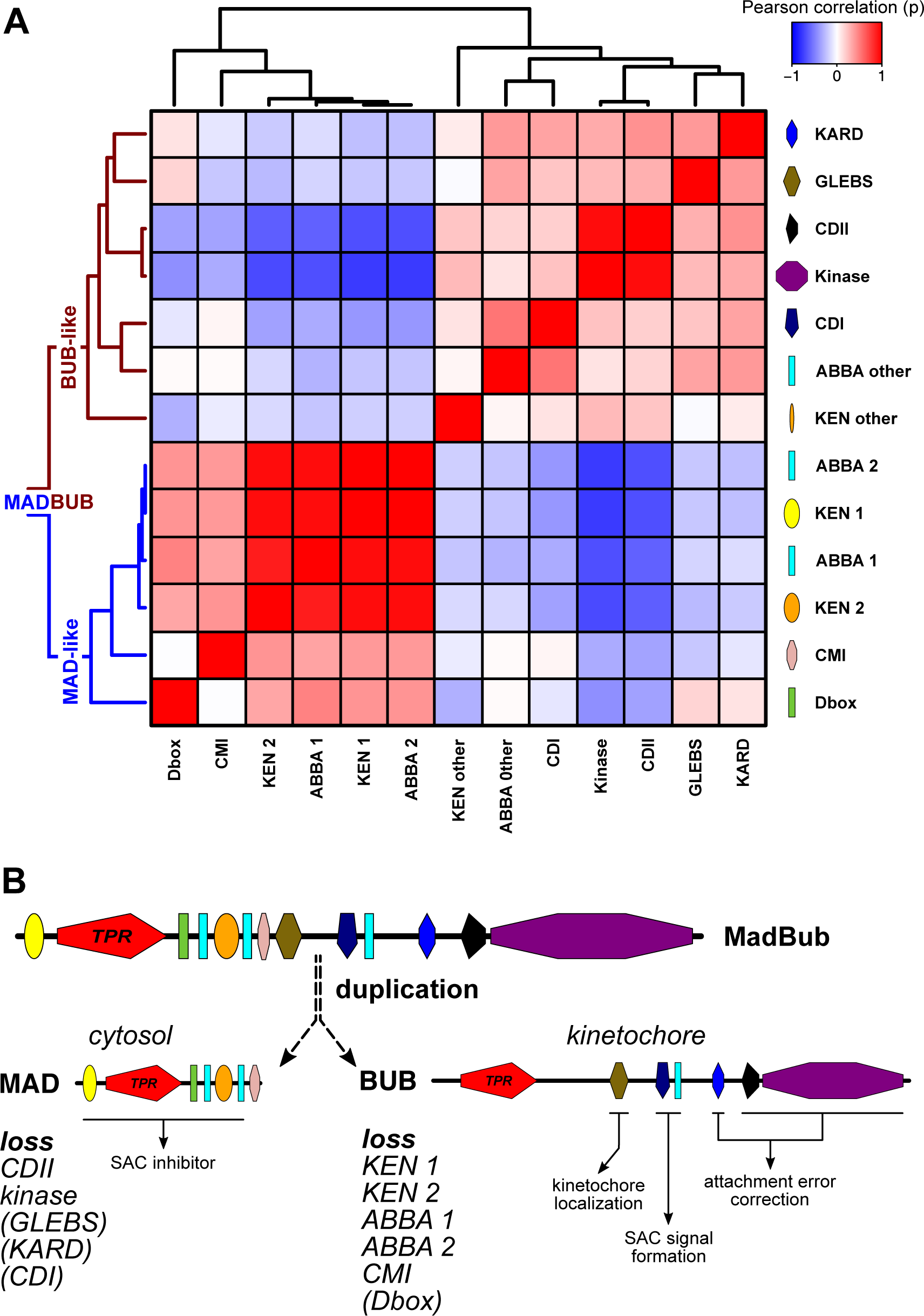
Co-evolution of conserved features signify subfunctionalization of MAD and BUB after MADBUB duplication. (A) Average clustering based on pearson distances of conserved ancestral feature correlation matrix (distance=1-r) of all MADBUB paralogs from figure 1. Red and blue indicate co-presence or absence of features in the same paralog, respectively. (B) Evolutionary scenario of MADBUB subfunctionalization: MAD (cytosol) as a SAC effector and BUB (kinetochore) involved in SAC signal formation and kinetochore microtubule attachment.

**Figure 3.**
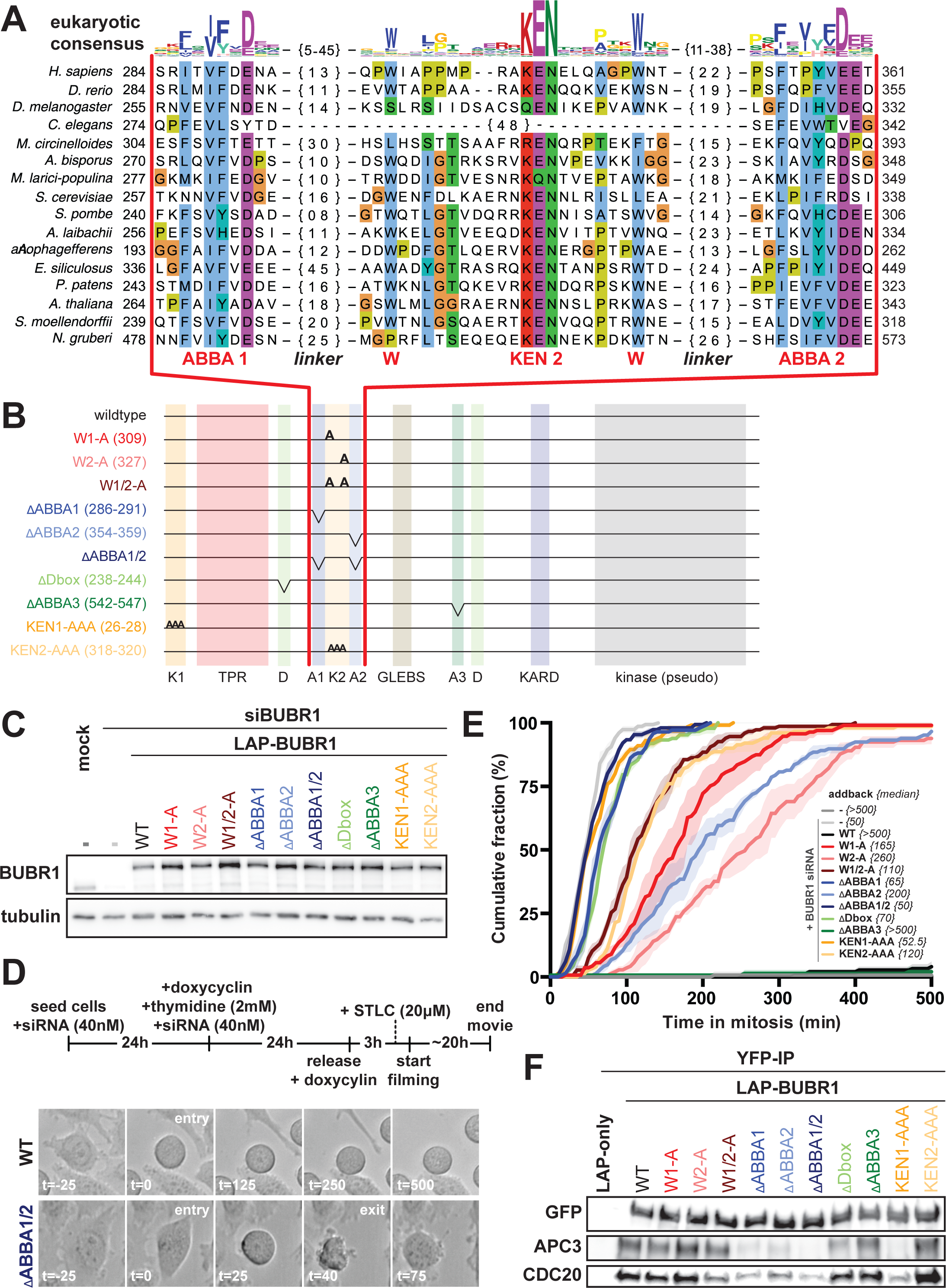
The evolutionary conserved cassette ABBA1-KEN2-ABBA2 in BUBR1 is essential for SAC signaling. (A) Alignment of ABBA1-KEN2-ABBA2 cassette (red). Linkers (black) between ABBA motifs and KEN2 are indicated by {n}. The sequence logo on top is representative for all eukaryotic sequences (colors reflect distinct amino acid properties and height of the letters indicates conservation of amino acids). (B) Schematic representation of LAP-hBUBR1 mutants. Color-coding is consistent for each mutant in this figure (C) Immunoblots of BUBR1 and tubulin of mitotic lysates of HeLa FlpIn cell lines stably expressing LAP-tagged BUBR1 proteins. Cells were treated with siRNA (40nM) for 48 hours and cells were released and arrested into taxol after double thymidine block. (E) Time-lapse analysis of HeLa FlpIn cells expressing hBUBR1 mutants, treated with 20 μM STLC. Data (n=50, N=3) indicate the mean of cumulative fraction of cells that exit mitosis after nuclear envelope breakdown. Transparent regions represent the standard error of the mean. Values between accolades {} indicate the median value. Cells were score by cell morphology using DIC imaging (D); see for examples of SAC deficient (ΔABBA1/2) and proficient cells (wildtype). Only YFP-positive cells were considered for analyses. (F) Immunoblots of GFP, APC3 and CDC20 in LAP-BUBR1 precipitations (LAP-pulldown) in whole cell lysates of mitotic HeLa FlpIn cells stably expressing LAP-BUBR1 mutant constructs.

### The conserved ABBA1-KEN2-ABBA2 cassette is essential for proper SAC signaling in human cells

The strong correlation of the ABBA1-KEN2-ABBA2 cassette with KEN1 and the D-box, urged us to examine the role of these motifs in BUBR1-dependent SAC signaling in human cells. We therefore generated stable isogenic HeLa FlpIn cell lines expressing doxycyclin-inducible versions of LAP-tagged BUBR1 (Suijkerbuijk et al., 2010). These included: ΔABBA1, ΔABBA2, ΔABBA1+2, alanine-substitutions of the two KEN2-flanking tryptophans (W1-A, W2-A and W1/2-A), KEN1-AAA KEN2-AAA, ΔABBA3 and ΔD-box (**figure 3a-c**). The SAC was severely compromised in cells depleted of endogenous BUBR1 by RNAi, as measured by inability to maintain mitotic arrest upon treatment with S-trityl-L-cysteine (STLC) (Ogo et al., 2007) (median(m) = 50 minutes (min.) from NEBD to mitotic exit, compared control (m>500 min.)) (**figure 3d-e**). SAC proficiency was restored by expression of siRNA-resistant LAP-BUBR1 (m>500 min.). As shown previously (Burton and Solomon, 2007a; Elowe et al., 2010; Lara-Gonzalez et al., 2011), mutants of KEN1, KEN2 and the D-box strongly affected the SAC. Importantly, BUBR1 lacking ABBA1 or ABBA2 or both, or either of the two tryptophans, could not rescue the SAC (**figure 3e**). We observed a consistently stronger phenotype for the mutated motifs on the N-terminal side of KEN2 (ΔABBA1 (m=65 min.) and W1-A (m=165 min.)) compared to those on the C-terminal side (ΔABBA2 (m=200 min.) and W2-A (m=260 min.)). The double ABBA (1/2) and tryptophan (1/2) mutants were however further compromised (m = 50 and 110 min., respectively), suggesting non-redundant functions. As expected from the interaction of ABBA motifs with the WD40 domain of CDC20 and CDH1 (DiFiore et al., 2015), BUBR1 lacking ABBA1 and/or ABBA2 was less efficient in binding APC/C-Cdc20 in mitotic human cells, to a similar extent as mutations in KEN1 (**figure 3f**). In our hands ABBA1 and 2 mutants were more strongly deficient in SAC signaling and APC/C-Cdc20 binding than the previously described ABBA motif (ABBA3) (**figure 3d-e**). In conclusion therefore, the ABBA1-KEN2-ABBA2 cassette in BUBR1 is essential for APC/C inhibition by the SAC.

We here discovered a symmetric cassette of SLiMs containing two Cdc20-binding ABBA motifs and KEN2. This cassette strongly co-occurs with KEN1 in MAD-like and MadBub proteins throughout eukaryotic evolution and has important contributions to the SAC in human cells. Our co-precipitation experiments along with the known roles for ABBA-like motifs and KEN2 and their recent modeling into the MCC-APC/C structure (Alfieri et al., 2016; Yamaguchi et al., 2016) strongly suggest that the ABBA1-W-KEN2-W-ABBA2 cassette interacts with one or multiple Cdc20 molecules. Together with KEN1, these interactions likely regulate affinity of MCC for APC/C or its positioning once bound to APC/C. The constellation of interactions between Cdc20 molecules and the various Cdc20-binding motifs in one molecule of BUBR1 (3x ABBA, 2x KEN and a D-box) is not immediately obvious, and will have to await detailed atomic insights. The symmetric arrangement of the cassette may be significant in this regard, as is the observation that (despite a highly conserved WD40 structure of Cdc20) the length of spacing between the ABBA motifs and KEN2 is highly variable between species. A more detailed understanding of SAC function may be aided by ConFeaX-driven discovery of lineage-specific conserved features in the MadBub family when more genome sequences become available, as well as of features in other SAC proteins families.

## Supplementary information

Supplementary information included Supplementary discussion, 2 figures, 3 tables and 2 sequence files.

## Methods

### Phylogenomic analysis

We performed iterated sensitive homology searches with jackhammer (Finn et al., 2011) (based on the TPR, kinase, CDI, GLEBS and KEN boxes) using a permissive E-value and bitscore cut-off to include diverged homologs on uniprot release 2016_08 and Ensemble Genomes 32 (http://www.ebi.ac.uk/Tools/hmmer/search/jackhmmer). Incompletely predicted genes were searched against whole genome shotgun contigs (wgs, http://www.ncbi.nlm.nih.gov/genbank/wgs) using tblastn. Significant hits were manually predicted using AUGUST (Stanke et al., 2006) and GENESCAN (Burge and Karlin, 1997). In total we used 152 MadBub homologs (**supplementary sequence file 1**). The TPR domains of 148 sequences were aligned using MAFFT-LINSI (Katoh and Standley, 2013); only columns with 80% occupancy were considered for further analysis. Phylogenetic analysis of the resulting multiple sequence alignment was performed using RAxML (Stamatakis, 2014) (**supplementary figure 1a**). Model selection was performed using Prot Test (Darriba et al., 2011) (Akaike Information Criterion): LG+G was chosen as the evolutionary model.

### Conserved Feature Extraction and subfunctionalization analysis

ConFeaX starts with a probabilistic search for short conserved regions (max 50) using the MEME algorithm (option: any number of repeats) (Bailey et al., 2009). Significant motif hits are extended on both sides by 5 residues to compensate for the strict treatment of alignment information by the MEME algorithm. Next, MAFFT-LINSI (Katoh and Standley, 2013) introduces gaps and the alignments are modeled using the HMMER package (Eddy, 2011) and used to search for hits that are missed by the MEME algorithm. Subsequent alignment and HMM searches were iterated until convergence. For short linear motifs with few conserved positions, specific optimization of the alignments and HMM models using permissive E-values/bit scores was needed (e.g. ABBA motif and D-box). Sequence logos were obtained using weblogo2 (Crooks et al., 2004). Subsequently, from each of the conserved features, a phylogenetic profile was derived (present is ‘1’ and absent is ‘0’) for all duplicated MadBub sequences as presented in **figure 1**. For all possible pairs, we determined the correlation using Pearson correlation coefficient (Wu et al., 2003). Average clustering based on Pearson distances (d=1-r) was used to indicate sub-functionalization.

### Cell culture, transfection and plasmids

HeLa FlpIn T-rex cells were grown in DMEM high glucose supplemented with 10% Tet-free FBS (Clontech), penicillin/streptomycin (50 mg ml^−1^), alanyl-glutamine (Sigma; 2 mM). pCDNA5-constructs were co-transfected with pOgg44 recombinase in a 10:1 ratio (Klebig et al., 2009) using FuGEHE HD (Roche) as a transfection reagent. After transfection, the medium was supplemented with puromycin (1 μg ml^−1^) and blasticidin (8 μg ml^−1^) until cells were fully confluent in a 10cm culture dish. siBUBR1 (5’-AGAUCCUGGCUAACUGUUCUU-3’ custom Dharmacon) was transfected using Hiperfect (Qiagen) at 40 nM for 48h according to manufacturer’s guidelines. RNAi-resistant LAP (YFP)-BUBR1 was sub cloned from plC58 (Suijkerbuijk et al., 2010) into pCDNA5.1-puro using AflII and BamHI restriction sites. To acquire mutants, site-directed mutagenesis was performed using the quickchange strategy (for primer sequences see **supplementary table III**).

### Live cell imaging

For live cell imaging experiments, the stable HeLa-FlpIn-TRex cells were transfected with 40nM siRNA (start and at 24 hrs). After 24 hrs, the medium was supplemented with thymidine (2.5 mM) and doxycyclin (2 μg ml^−1^) for 24 hrs to arrest cells in early S-phase and to induce expression of the stably integrated construct, respectively. After 48 hrs, cells were released for 3 hrs and arrested in prometaphase of the mitotic cell cycle (after ~8-10 hrs) by the addition of the Eg5 inhibitor S-trityl-L-cysteine (STLC, 20 μM). HeLa cells were imaged (DIC) in a heated chamber (37 °C, 5% CO_2_) using a CFI S Plan Fluor ELWD 20x/NA 0.45 dry objective on a Nikon Ti-Eclipse wide field microscope controlled by NIS software (Nikon). Images were acquired using an Andor Zyla 4.2 sCMOS camera and processed using NIS software (Nikon) and ImageJ.

### Immuno-precipitation and Western blot

HeLa-FlpIn-TRex cells were induced with doxycyclin (2 μg ml^−1^) 48 hrs before harvesting. Synchronization by thymidine (2 mM) for 24 hrs and release for 10 hrs into Taxol (2 μM) arrested cells in prometaphase. Cells were collected by mitotic shake-off. Lysis was done in 50 mM Tris-HCl (pH 7.5), 100 mM NaCl, 0.5% NP40, 1mM EDTA, 1mM DTT, protease inhibitor cocktail (Roche) and phosphatase inhibitor cocktails 2 and 3 (Sigma). Complexes were purified using GFP-Trap beads (ChromoTek) for 15 min at 4°C. Precipitated proteins were washed with lysis buffer and eluted in 5x SDS sample buffer. Primary antibodies were used at the following dilutions for western blotting: BUBR1 (A300-386A Bethyl) 1:2000, alpha-Tubulin (T9026 Sigma) 1:5000, GFP (Custom) 1:10 000, APC1 (A301-653A Bethyl) 1:2500, APC3 (gift from Phil Hieter) 1:2000, MAD2 (Custom) 1:2000, CDC20 (A301-180A Bethyl) 1:1000. Western blot signals were detected by chemiluminescence using an ImageQuant LAS 4000 (GE Healthcare) imager.

## Author contributions

ET performed the motif search, phylogenetic analysis and SAC assays. DB performed the immunoprecipitation and western blot analyses. GK, BS conceived and managed the project. ET, BS and GK wrote the manuscript.

## Acknowledgements

The authors thank the Snel and Kops lab for discussion and feedback. We thank Bas de Wolf and Laura Demmers for making cell lines. This work was supported by the UMC Utrecht and is part of the VICI research programme with project number 016.160.638, which is (partly) financed by the Netherlands Organisation for Scientific Research (NWO). DB was supported by a DFG fellowship with number BA 5417/1-1, which is financed by the German Research Foundation (DFG).

